# Spatio-temporal development of the urban heat island in a socioeconomically diverse tropical city

**DOI:** 10.1101/2022.07.07.499124

**Authors:** Emma E. Ramsay, Grant A. Duffy, Kerrie Burge, Ruzka R. Taruc, Genie M. Fleming, Peter A. Faber, Steven L. Chown

**Affiliations:** School of Biological Sciences, Monash University, Victoria, 3800, Australia; Department of Marine Science, University of Otago, Dunedin, New Zealand; Monash Sustainable Development Institute, Monash University, Victoria 3800, Australia; RISE Program, Faculty of Public Health, Makassar, Hasanuddin University, Makassar, Indonesia

**Keywords:** urban heat islands, informal settlements, tropics, urbanisation, cities

## Abstract

Urban heat islands, where temperatures are elevated relative to non-urban surrounds, are near-ubiquitous in cities globally. Yet, the magnitude and form of urban heat islands in the tropics, where heat has a large morbidity and mortality burden, is less well understood, especially for socioeconomically diverse communities such as those living in urban informal settlements. We utilised 29 years of Landsat satellite-derived surface temperature, corroborated by *in situ* measurements, to provide a detailed spatial and temporal assessment of urban heat islands in Makassar, Indonesia, a city that is representative of rapidly growing urban settlements across the tropics. We did so with explicit consideration of vulnerable communities living informally. Our analysis identified surface urban heat islands of up to 9.2 °C in long-urbanised parts of the city and 6.3 °C in informal settlements, the seasonal patterns of which were driven by change in non-urban areas rather than in urban areas themselves. In recently urbanised areas, the majority of urban heat island increase occurred before areas became 50% urbanised. As tropical cities continue to expand we expect that urban heat islands will develop quickly as land is urbanised, whereas the established heat island in long-urbanised areas will remain stable in response to city expansion. Green and blue space protect some informal settlements from the worst urban heat islands and maintenance of such space will be essential to mitigate the growing heat burden from urban expansion and anthropogenic climate change. We advocate for green space to be prioritised in urban planning, redevelopment and informal settlement upgrading programs, with consideration of the unique environmental and socioeconomic context of tropical cities.

**Highlights:**

- Long-term, fine-scale data are essential to understand the dynamics of urban heat
- Surface heat islands reached 9.2 °C in the urban core, 6.3 °C in informal settlements
- *In situ* data support the use of remote sensing for heat island characterisation
- The majority of heat island growth occurred before land was 50% urbanised
- Green and blue space can mitigate heat in informal settlements

## 1. Introduction

Human populations are rapidly expanding and becoming increasingly urbanised. More than half of the world’s population now lives in urban areas and this is expected to increase to nearly 70% by 2050 (United Nations 2019), accompanied by a near doubling of global urban land cover between 2015 and 2050 (Huang *et al*. 2019). The majority of this growth will occur in developing countries, particularly in Asia and Africa, home to most of the world’s fastest growing cities (Laurance *et al*. 2015; United Nations 2019). City growth in these contexts is often characterised by urban informal settlements which typically have poor quality infrastructure and services and high exposure to environmental hazards (Ezeh *et al*. 2017). More than one billion people live in informal settlements globally, primarily in East and South-East Asia, a number which is expected to increase with continued population growth and urbanisation (UN-Habitat 2015; United Nations 2021b).

The growth of cities drives local change in land cover, climate and hydrological cycles (Grimm *et al*. 2008). One of the most prominent outcomes of this environmental change is the urban heat island (UHI) effect (Seto & Shepherd 2009; Bai *et al*. 2017). Urban heat islands are primarily caused by land cover changes, associated with urbanisation, which alter the surface energy balance and morphology (Oke 1982; Arnfield 2003). Such land cover change across a broad extent leads to high absorption and retention of radiant heat in urban areas, and thus increased ambient and surface temperature relative to surrounding non-urban areas (Bai *et al*. 2017). Elevated temperatures from UHIs can exacerbate extreme heat (Founda & Santamouris 2017; Zhao *et al*. 2018), which has adverse impacts on human health and wellbeing (Tan *et al*. 2010; Ebi *et al*. 2021a). Througout the tropics, UHIs compound year-round high temperature and humidity, which is of particuclar concern for vulnerable populations such as those living in urban informal settlements (Scott *et al*. 2017; Ramsay *et al*. 2021) who have limited capacity to adapt (Pasquini *et al*. 2020) and a large exisiting health burden (Ezeh *et al*. 2017; Lilford *et al*. 2017).

Urban heat islands have been relatively well characterised globally (Peng *et al*. 2012; Chakraborty & Lee 2019), with advances in remote sensing techniques and data availability accelerating this characterisation (Kotharkar *et al*. 2018). Much of the focus remains, however, on temperate regions of the world including Europe, North America and China (Zhou *et al*. 2018). Many studies are also limited to simplified comparisons of urban and non-urban areas, or single time-points, which do not sufficiently capture intra-urban variation, temporal trends or local context (Tuholske *et al*. 2021). Spatial and temporal patterns of urban warming are heterogenous globally. UHIs are influenced by local climatic conditions, city size and density (Imhoff *et al*. 2010; Li *et al*. 2017; Miles & Esau 2020), and variation in urban land use, morphology and socioeconomics (Chen *et al*. 2006; Wang *et al*. 2019). Evidence is mounting that in some cities lower income neighbourhoods have disproportionately high exposure to UHIs (Buyantuyev & Wu 2009; Chakraborty *et al*. 2019). Such neighbourhoods are likely to include informal settlements, which themselves may be exposed to higher UHIs due to dense housing and limited green space (Mehrotra *et al*. 2018; Wang *et al*. 2019), although this may be reduced if settlements are embedded in green or blue space (Jacobs *et al*. 2019). However, spatially explicit information about informal settlements is scarce (Satterthwaite *et al*. 2020) and they have thus been insufficiently considered in UHI analyses.

In the face of expected growth in informal settlement extent and numbers of residents, coupled with the rising health and socioeconomic burdens associated with lack of services, climate change and their interactions (Scovronick *et al*. 2015; Satterthwaite *et al*. 2020), improving these settlements is a growing international priority. This priority is reflected in the increasing interest and investment by development banks in nature-based solutions (RISE & ADB 2021; Hamel & Tan 2022) which are a major focus of both the Sustainable Development Goals (United Nations 2021b) and Planetary Health approaches (Whitmee *et al*. 2015). Mitigating the extent and impacts of UHIs must clearly form an essential component of improving informal settlement environments, and is potentially most feasible where nature-based solutions are deployed. The basis for such mitigation is the development of an explicit, localised understanding of the spatial and temporal patterns of UHI exposure, and the likely impacts thereof in rapidly growing cities and their associated informal settlements (Deilami *et al*. 2018; Nagendra *et al*. 2018). By understanding trends in urban development and UHIs, the potential for spatial expansion and intensification of UHIs with continued urbanisation can be projected (Li *et al*. 2021). In turn, this understanding of spatiotemporal trends can be integrated into context-specific mitigation strategies.

Here we quantify spatial and temporal patterns of urbanisation and accompanying UHIs over the last 30 years in an exemplar, socioeconomically diverse, tropical South-East Asian city: Makassar, South Sulawesi, Indonesia. Makassar is a medium sized (< 5 million people) city with a population of ~1.5 million people, in this way representing more than 90% of urban settlements globally (United Nations 2018). The city is currently characterised by a diversity of informal settlements distributed across much of its extent (French *et al*. 2021). Indonesia is one of the most urbanised countries in Asia with 110 million people living in 60 cities (Gunawan *et al*. 2015) and 30% of the urban population living informally (United Nations 2021a). Our aim is to provide improved understanding of spatio-temporal trends of UHIs and identify the likely drivers of these trends as the city has grown, as an exemplar of how UHIs develop in such settings and in particular in informal settlements, with a view to integrating mitigation in city development and upgrading, including the nature-based informal settlement revitalisation proposed both for this city and for others (Leder *et al*. 2021).

## 2. Materials and methods

Local scale UHI analyses require data at fine spatial and temporal resolutions. Ambient temperature is preferred for analysing urban heat in terms of impacts on humans, yet climate data from traditional weather stations do not capture localised heat exposure in complex urban settlements (Scott *et al*. 2017; Ramsay *et al*. 2021) and are particularly scarce throughout the tropics (Zaitchik & Tuholske 2021). Satellite data are therefore valuable for fine scale spatial and temporal analyses where *in situ* data are unavailable (Zhou *et al*. 2018), and have thus been used extensively to characterise UHIs across the globe (eg. Peng *et al*. 2012; Estoque *et al*. 2017; Zhou *et al*. 2017; Chakraborty & Lee 2019; Manoli *et al*. 2019).

We therefore used the 40 year near-global record of multispectral spatial data from the Landsat satellite series, including Landsat 5 TM, Landsat 7 ETM+ and Landsat 8 OLI TIR between 1980 and 2020 (Wulder *et al*. 2019) to characterise urban land cover and analyse spatial and temporal trends in surface temperature in Makassar, Indonesia. Landsat data comprise spectral bands in the visible, near infrared, short wave and thermal infrared spectra at a spatial resolution of 30 m (thermal bands are resampled to 30 m), with a 16-day temporal coverage of the earth’s surface (Markham & Helder 2012; Roy *et al*. 2014). Landsat satellites have an overpass time of approximately 10 am Makassar local time (WITA; UTC+8).

### 2. 1 Urban land cover change

Supervised land cover classifications were performed by E.E.R. at 11 time points between 1993 and 2019 to determine changes in urban land cover over time (Figure 1). Landsat Collection 1, Level 1 imagery (U.S. Geological Survey 2019) were downloaded and pre-processed (conversion to reflectance, DOS1 atmospheric correction) using the Semi-Automatic Classification Plugin (version 7.10.5; Congedo 2016) in QGIS (version 3.4.2; QGIS Development Team 2018). Images within the dry season months (April to October) were manually selected for land cover classification to minimise seasonal variation in land cover and optimise availability of cloud-free images. Where data from more than one satellite were available, data from the most recently deployed satellite were used, except between 2003 and 2013, where Landsat 5 were used in place of Landsat 7 after the failure of the scan line corrector on Landsat 7 (Markham *et al*. 2004).

**Figure 1.**
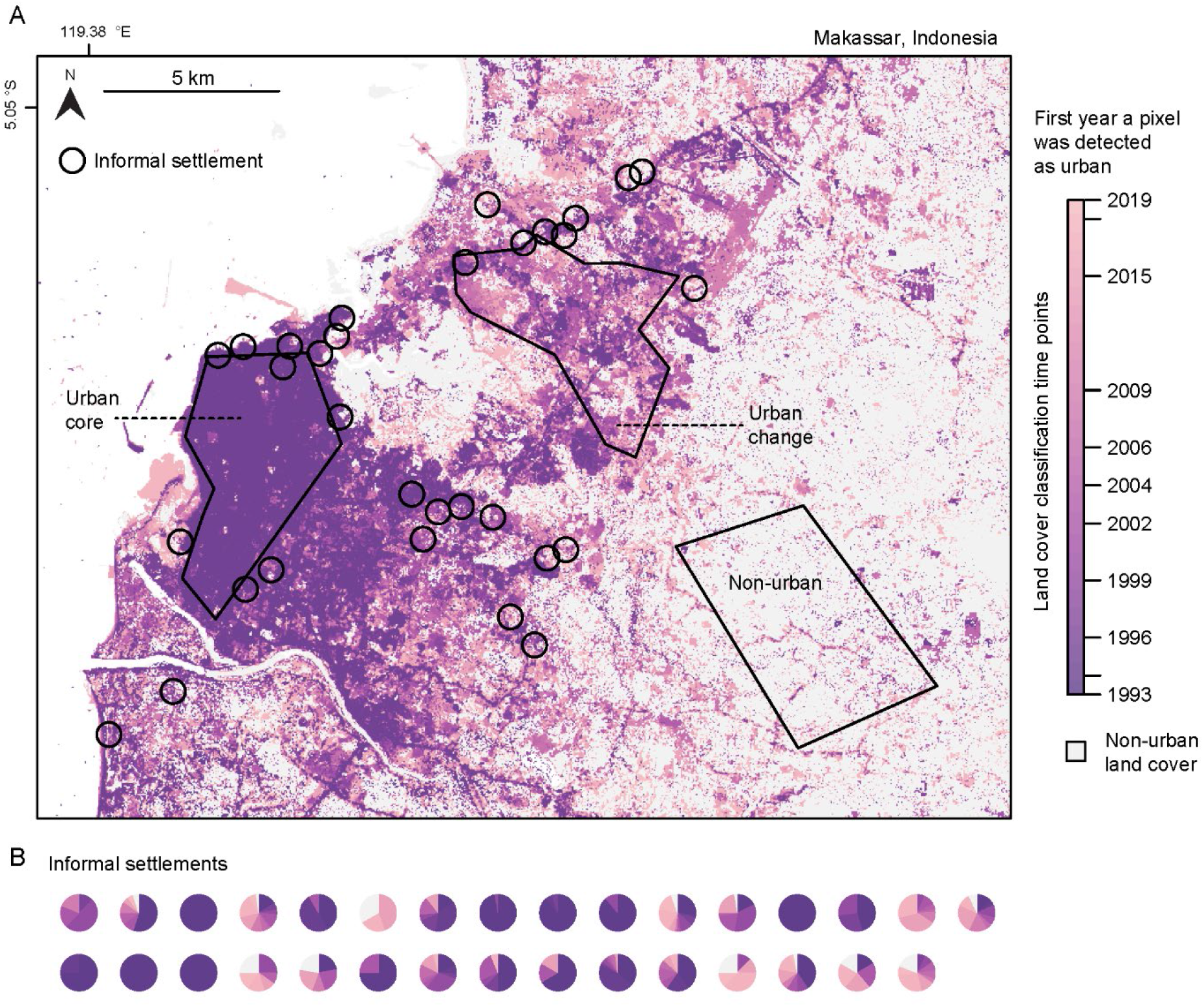
A) Urban land cover change in Makassar, Indonesia between 1993 and 2019 showing the urban core, urban change, non-urban and informal settlement sampling areas. Informal settlement locations represent the general location of the settlement rather than the exact polygon. Map shows the first year a pixel was detected as urban based on 11 land cover classifications. Legend ticks represent land cover classification time-points. B) The distribution of urban land cover change in informal settlements. Each pie chart represents the first year a pixel was detected as urban in one settlement

The Semi-Automatic Classification Plugin (Congedo 2016) was used to classify land cover at each time point based on remote-sensed visible spectrum and infrared Landsat bands (Landsat 5 and Landsat 7: Bands 1-5 and 7, Landsat 8: Bands 2-7) using the maximum likelihood classification method. Clouds were masked manually by drawing polygons over areas affected by clouds or their shadows and then images were clipped to a study region which included the urban extent of the city as well as a variety of non-urban land cover types for training. The classification algorithm was trained by manually selecting training areas from true and false colour composites of Landsat bands and assigning them to six broad land cover classes: water, urbanised, vegetation, cropland, wet-cropland/wetland and bare soil, based on visual assessment and photointerpretation of the Landsat images. Training areas were a minimum of 60 pixels and a maximum of 1000 pixels. A minimum of 10 training areas were selected for each land cover class and these were spread throughout the image to obtain a representative description of each class (Foody & Mathur 2004).

Based on changes in urbanised land cover over time (Figure 1A) we defined four spatial regions for UHI analysis (Figure 1A):

a. The urban core which was predominantly urbanised at the first land cover classification (23 June 1993) and remained urbanised over the time period of this study
b. An area of urban change which changed from predominantly non-urban to urbanised over the time period of this study
c. Informal settlements (detailed below)
d. A non-urban area for comparison which comprised primarily non-urban land cover types and remained predominantly non-urban for the entire time period of this study

We manually defined the urban core, urban change and non-urban areas based on the land cover classifications (Figure 1A), whilst minimising geographical and topographical differences which could confound temperature observations. Informal settlement locations in Makassar were identified from a list of candidate settlements for the Revitalising Informal Settlements and their Environments (RISE) Program (Leder *et al*. 2021). These settlements were visited and verified as informal settlements by K.B. The initial list included 39 informal settlement locations (including the 12 settlements in the RISE Program and the RISE demonstration site; RISE & ADB 2021). Where settlements were less than 200 m from each other we grouped them as a single spatial unit, resulting in a final dataset of 31 settlements. The settlements are geographically dispersed across the city (Figure 1A) and represent some of the most vulnerable urban dwellers (French *et al*. 2021). Settlements have a mean area of 0.019 km^2^, which is comparable to the global average of 0.016 km^2^ (Friesen *et al*. 2018).

### 2.2 Land surface temperature

From the Landsat satellite archive (Cook *et al*. 2014), we downloaded all available Landsat Collection 2, Level 2 imagery (U.S. Geological Survey 2020) (aside from Landsat 7 after 2003) for Makassar (path 114, row 64) between 1970 and 2020 using the Earth Explorer Bulk Download Application (https://earthexplorer.usgs.gov/bulk/), totalling 376 time points. Surface temperature and pixel quality rasters were extracted for each time point, scaled and converted from Kelvin to degrees Celsius (U.S. Geological Survey 2020). Surface temperature rasters were masked using the pixel quality raster by setting pixels identified as clouds or water to NA. We also extracted the red (R) and near-infrared bands (NIR) to calculate the Normalised Difference Vegetation Index (NDVI) as:

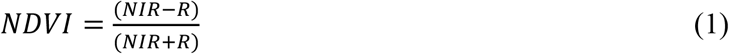

We randomly sampled 31 points within each of the spatial regions (urban core, urban change and non-urban; Figure 1) and then buffered them (by a radius of 78m or ~2.6 pixels) to approximately the same size as informal settlements (~21 pixels). This enabled us to achieve a balanced sampling design with the informal settlement locations and minimised the effect of spatial autocorrelation, by analysing a random sample of surface temperature observations rather than all pixels within each region (Buyantuyev & Wu 2009).

Landsat satellite data is limited by cloud cover, especially during the wet season (November to March) where few cloud free images are available. To maximise data availability for analyses we filtered surface temperature data in two stages. Time points with more than 25% missing data in the general study area (map area in Figure 1A) and more than 10% of missing data in the sampling areas (Figure 1A) were excluded. The remaining data were manually checked for cloud contamination not sufficiently masked by the pixel quality raster, by visualising the surface temperature rasters, resulting in 89 time points between 1991 and 2020 available to be included in analyses. Data were filtered for a second time for each set of analyses, based on the data required for each model. We excluded random sample patches with any NA pixels (indicating that a cloud was close by and may have affected surface temperature data) and excluded time points where there were more than six missing samples from the 31 patches or informal settlements.

Mean surface temperature and NDVI values were extracted for each sampling location (31 informal settlements and 31 randomly sampled patches in each of urban core, urban change and non-urban; Figure 1) using the *extract* function in the *raster* package (Hijmans 2020). Land cover from the 11 land cover classifications was also extracted for each and the percentage of urban pixels within the patch computed for each classification. The mean elevation, extracted from the Shuttle Radar Topography Mission (Farr *et al*. 2007) was between 0-30 m for all settlements and random patches.

Unless otherwise stated, all data processing was performed in R (version 4.05; R Core Team 2021) using the *dplyr* (Wickham *et al*. 2022), *raster* (Hijmans 2020), *sf* (Pebesma 2018) and *rgeos* (Bivand & Rundel 2020) packages.

### 2.3 Statistical analyses

Generalised additive mixed models (GAMMs) were used to estimate spatial and temporal trends in surface temperature. Additive models can fit smooth non-linear trends (such as trends over time) and can handle irregular temporal spacing of samples (Simpson 2018), thus making them suitable for our data. Random effects were incorporated to account for spatial autocorrelation. GAMMs were fit in R using the *mgcv* package (Wood 2004) with random effects estimated in the *nlme* package (Pinheiro *et al*. 2021) using restricted maximum likelihood estimation (REML; Wood 2011). Unless otherwise stated smooth trends were fit with thin plate regression splines (Wood 2003). Model selection was based on Akaike’s information criterion (AIC).

First, we modelled long-term trends (1991-2020) in surface temperature in the urban core, urban change and non-urban areas to understand if the magnitude of UHIs has changed over time with the growth of the city. Second, we modelled surface temperature in the urban change area separately to understand how change in urban land cover has driven changes in land surface temperature. Third, we quantified and compared current (2017-2020) UHIs in the urban core and informal settlements. Finally, we modelled spatial predictors of surface temperature in informal settlements. We visualised temporal trends and quantified UHIs from fitted data with simultaneous confidence intervals computed by simulating from the posterior distribution of the model using the *Gratia* package (Simpson 2018; Simpson 2022).

#### 2.3.1 Long-term trends

We modelled surface temperature in the urban core and non-urban areas as smooth functions of the long-term trend as days since first sampling date (with the first sampling date being day one) and the seasonal (within year) pattern modelled as a cubic spline smooth of day of year. Each smooth trend was fit separately for urban and non-urban data, which lowered AIC compared to models with one trend. Spatial plots of the normalised residuals indicated residual spatial autocorrelation in the model so we added a residual correlation structure, nested within each time point, which improved both visual inspection of plots and AIC. The optimal spatial correlation structure available in *nlme* was selected via AIC.

We fit the same model as above for surface temperature in the urban change and non-urban areas, with the addition of a smooth (tensor product) interaction between days since first sampling date and day of year to allow for the seasonal trend changing over time with land cover change.

#### 2.3.2 Urban change

Urbanisation occurred at different times across the urban change area (Figure 1). We therefore wanted to model the temperature trend separately for each randomly sampled patch to understand how UHIs develop as land converts to urban. We fit models following Pedersen *et al*. (2019) which allowed long-term smooths to be estimated for each urban change patch. The best model included a global smooth for the overall long-term trend and patch-level smooths modelled as a *t2* factor interactions which allowed the shape of the smooth to vary for each patch (Wood *et al*. 2012). The model included a seasonal smooth and a residual spatial correlation structure as detailed above.

We identified significant periods of change in surface temperature by computing the first derivative of the estimated long-term trends for each patch using the method of finite differences, where the difference between two very close together values is an approximation of the first derivative (Simpson 2018). We calculated the yearly mean derivative and compared these to the percentage of urban land cover in that year (interpolated linearly between land cover classification dates). Statistically significant periods of change were inferred where simultaneous confidence intervals of the derivative did not include zero (Simpson 2018).

Data exploration revealed very different shaped trends which seemed to depend on different urban land use in the area of urban change, which is dominated by more industrial areas compared to the urban core (Surya *et al*. 2021). To differentiate between different land use types we calculated the mean area of building footprints (extracted from Open Street Map; OpenStreetMap contributors 2022) intersecting each urban change patch. We classified patches as industrial land use where the mean building footprint size was greater than 250 m^2^ or as other urban land use when less than 250 m^2^. We then manually checked each patch using Google Earth Imagery and reclassified one patch as industrial where large buildings visible on Google Earth Imagery were missing from Open Street Map (see Figure S1 for examples of each land use type).

#### 2.3.3 Informal settlements

To quantify UHIs in the urban core and informal settlements we modelled recent (2017 - 2020) surface temperature in the urban core, informal settlements and non-urban areas. As in previous models, the seasonal trend was modelled as a smooth cubic spline of day of year, separately for each group, and a residual spatial correlation structure was included, nested within each time point. Seasonal patterns in UHIs (ΔT) were quantified by computing differences between fitted trends (Simpson 2022). Significant differences in temperature and thus UHIs were inferred where the simultaneous confidence interval of the difference did not include zero (Rose *et al*. 2012).

To examine spatial variation in UHI magnitude among informal settlements we modelled surface temperature as a smooth function of the mean NDVI of the settlement, the mean NDVI within a 250 m radius surrounding the settlement, and the distance to the coast. As in previous models, the seasonal trend was modelled as a smooth cubic spline of day of year, separately for each group and a residual spatial correlation structure was included, nested within each time point. We fit the same model with spatial predictors as linear parametric terms and only day of year as a smooth function.

### 2.4 In situ temperature validation

To understand the extent to which surface temperature is representative of conditions experienced by people living in informal settlements we compared satellite derived surface temperature to *in situ* temperature measurements in a subset of 12 informal settlements. Monitoring was undertaken as a part of the RISE Program (Leder *et al*. 2021) and is detailed in Ramsay *et al*. (2021). Hourly temperature was measured by a network of iButton data loggers (Maxim Integrated, San Jose, California), with approximately 60 loggers in houses (six in each of ten houses) and five outdoors in each settlement. Data retrieval was limited by logger loss, logger failure and fieldwork limitations over the Covid-19 pandemic. To capture representative conditions in each settlement we only included time periods where data were retrieved from at least two outdoor loggers at a settlement or loggers in at least two houses in a settlement (Table S1).

*In situ* temperature data that met the above requirements overlapped with eight satellite overpass days (Table S1). For each settlement we calculated daily mean, minimum and maximum ambient temperature, separately for indoors and outdoors, and compared these to the mean surface temperature of the settlement on the same day. Minimum and maximum temperatures were derived by calculating the minimum (maximum) daily temperature for each logger and then taking the mean of the minimum (maximum) values across each settlement.

We compared mean settlement surface temperature with *in situ* derived variables via ranged major axis regression using the R package *lmodel2* (Legendre 2018). Ranged major axis regression (or model II regression) aims to describe the relationship between *x* and *y* where both variables are not controlled by the researcher, and have natural variation and measurement error (Warton *et al*. 2006; Sokal & Rohlf 2012). We assumed that the error variance was larger for surface temperature, but proportional to the variance, thus making ranged major axis regression a suitable method for our data. The significance of the ranged major axis slope was tested via permutation (n permutations = 999).

## 3. Results

### 3.1 Urban land cover change

In Makassar, urban land cover increased by 175% (annual growth rate of 6.7%), from 65 km^2^ in 1993 to 179 km^2^ in 2019, across the map area in Figure 1A. Urban growth occurred primarily to the North-East of the city, where we examined associated change in UHIs. In informal settlements, urban land cover was mixed between recent and long urbanised land (Figure 1B). Overall, informal settlements comprised between 42.8% and 100% urbanised land cover based on the most recent land classifications (2018 and 2019).

### 3.2 Urban heat islands

#### 3.2.1 Urban core

The urban core, which represents long-urbanised parts of the city (Figure 1A), experienced significant UHIs across a typical year (modelled with data between 2017 and 2020), with a strong seasonal dependence (Figure 2; Table S2). The UHI averaged 4.5 °C across the year, reaching as high as 9.2 °C in February, but becoming a “cool island” by as much as −6.8 °C in October (Figure 2). Importantly, this seasonal pattern was due to elevated temperature in the non-urban area rather than a decrease in temperature in the urban core (Figure 2A). Such seasonal trends are consistent across our analyses where surface temperature in urban areas varies little seasonally, but seasonal changes in UHIs are driven by variation in non-urban areas. UHIs in the urban core have remained relatively stable over the last 30 years, with long-term trends in surface temperature showing little change (Figure 3A).

**Figure 2.**
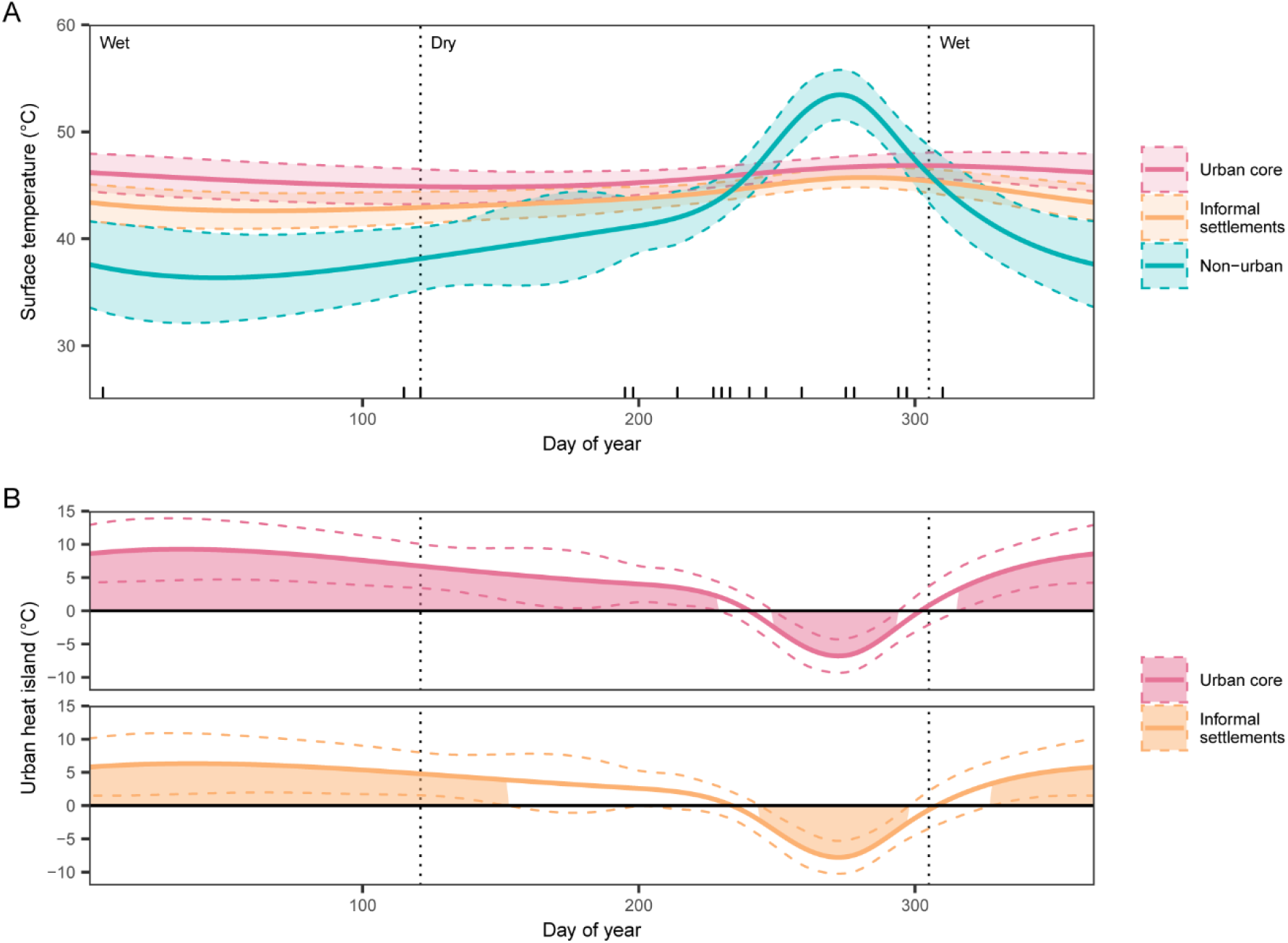
A) Fitted seasonal trends in surface temperature (2017-2020) with 95% confidence intervals. Ticks above X-axis represent sampling time points. B) Urban heat islands (ΔT) in the urban core and informal settlements. Shading above or below the fitted line indicates significant differences in surface temperature.

**Figure 3.**
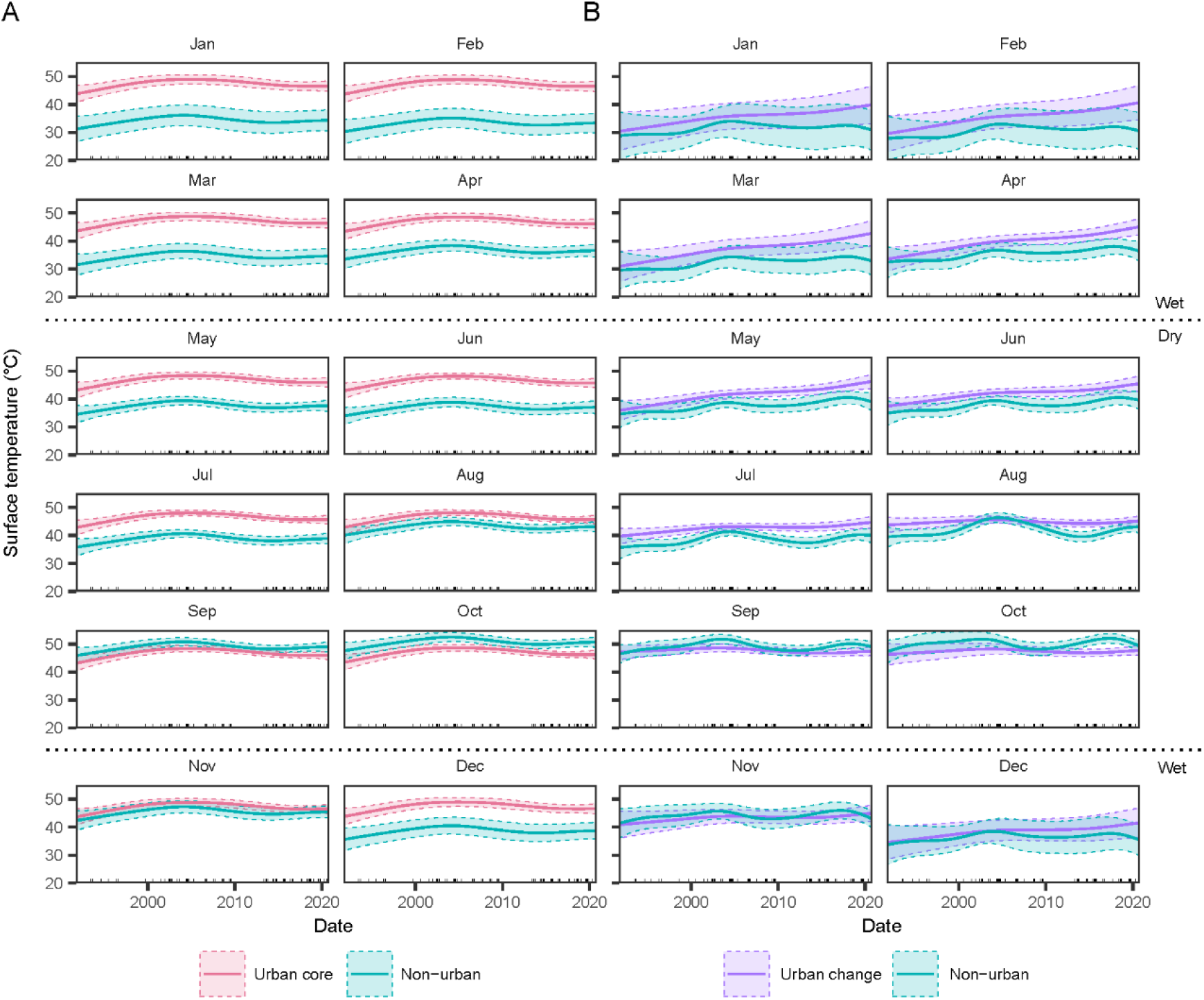
Fitted long-term trends in surface temperature for each month with 95% confidence intervals in A) the urban core and non-urban areas and B) urban change and non-urban areas. Fitted trends were computed for the 15^th^ day of each month. Ticks above the x-axis represent sampling time points included in the model.

#### 3.2.2 Urban change

Modelled long-term trends in surface temperature in the area of urban change reveal how UHIs have grown with urbanisation (Figure 3B, Figure 4). Trends differ by month as the GAMM allowed the seasonal pattern to vary with the long-term trend as land cover changed (Figure 3B; Table S2). Overall, surface temperature increased over the last 30 years, particularly between January and June where UHIs are largest in the urban core (Figure 3). In February, for example, the UHI increased from 1.7 °C in 1991 to 10.2 °C in 2020 (Figure 3B).

**Figure 4.**
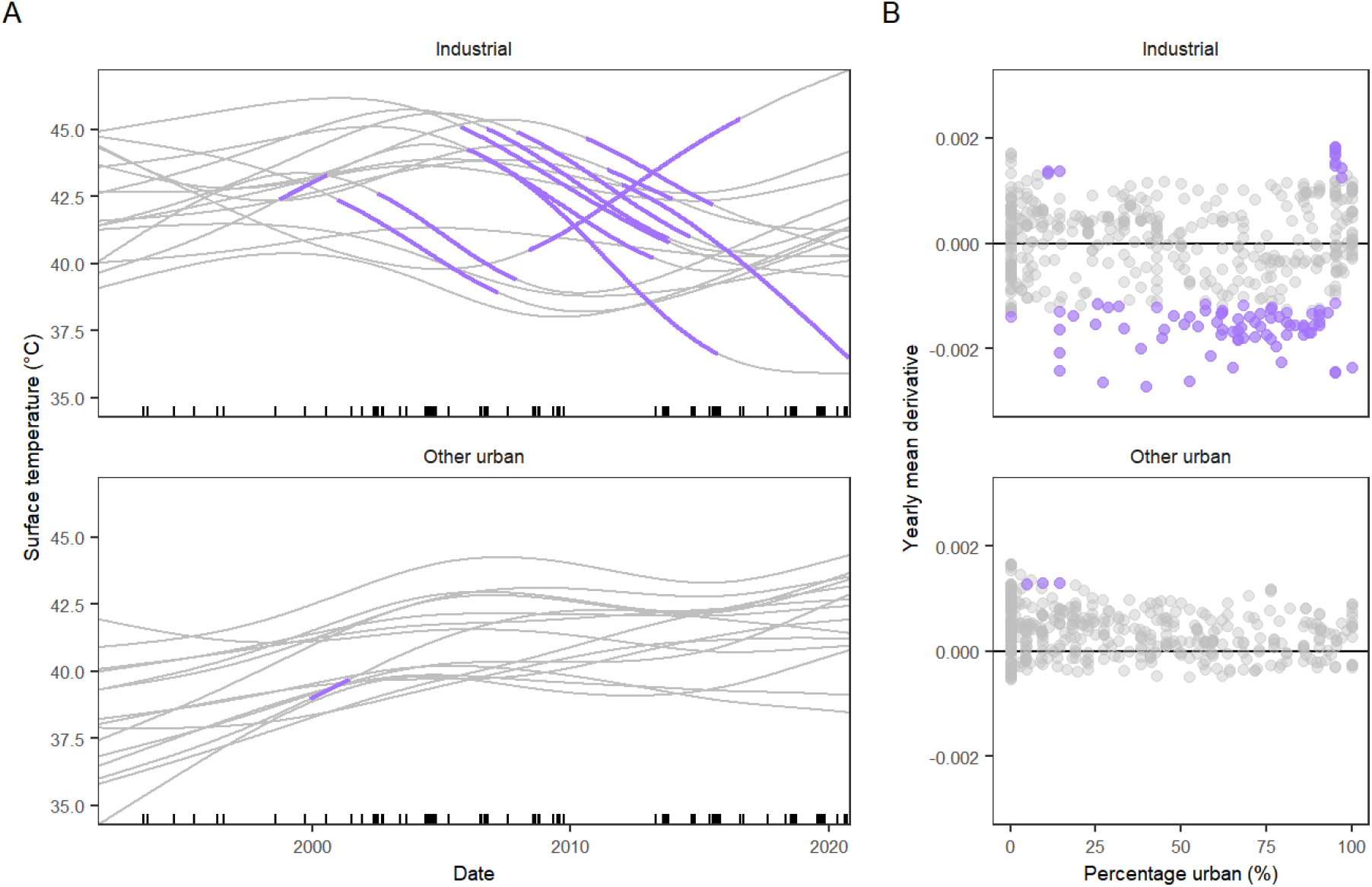
A) Fitted long-term trends in surface temperature in randomly sampled urban change patches, split by industrial and other urban areas. Fitted trends were computed on the 183^rd^ day of the year. Ticks above the x-axis represent sampling time points included in the model. B) Yearly mean derivative of the long-term trend and the percentage urban land cover of the patch in that year. Significant periods of change, identified from the derivative of the trend, are highlighted in purple.

Long-term trends modelled for each randomly sampled patch (Table S2) were much more mixed, accounting for the wide confidence intervals in Figure 3B. The area of urban change is dominated by industrial land use which is not typical of the urban core (Surya *et al*. 2021). Trends in industrial areas (mean building footprint > 250 m^2^), were mixed with periods of both significant increase and decrease, identified from the derivative of the trend (Figure 4). Trends in other urban areas (mean building footprint < 250 m^2^), which encompass suburban areas, were much more consistent, showing only periods of significant increase (Figure 4). Across other urban patches, 73% of years where surface temperature increased (positive derivative) occurred before a patch was 50% urbanised, indicating that UHIs establish early on as urban land is converted (Figure 4b).

#### 3.3.3 Informal settlements

UHIs in informal settlements followed a similar seasonal pattern to the urban core but were on average 1.9 °C lower (Figure 2). UHIs still exceeded 4 °C (averaging 2.6 °C) for much of the year and reached as high as 6.3 °C in February and as low as −7.8 °C in October. The average NDVI of settlements, and surrounding settlements, as well as distance to the coast were important predictors of surface temperature among informal settlements (Figure 5; Table S2), explaining nearly 40 % of the deviance (adj-R^2^ = 0.385). Settlements further from the coast had higher surface temperatures, with larger warming effects when further than 4 km and larger cooling effects when within 1 km of the coast (Figure 5). The NDVI of a settlement had the largest effect, where settlements with NDVI less than 0.2 were considerably warmer (effect of more than 5 °C) than those with NDVI greater than 0.2. For comparison, the mean NDVI of randomly sampled patches in the urban core was 0.22 between 2017 and 2020 (ranging between 0.20 in October and 0.26 in January). The NDVI of the 250 m surrounding each settlement had a smaller effect but settlements surrounded by land with higher NDVI, indicating more green space, were cooler. The inclusion of NDVI as a predictor negated the seasonal effect (Figure 5), indicating that seasonal changes in vegetation are responsible for much of the seasonal variation in surface temperature. The direction of effects was consistent when predictors were modelled as linear parametric terms (Table S3; adj-R^2^ = 0.411).

**Figure 5.**
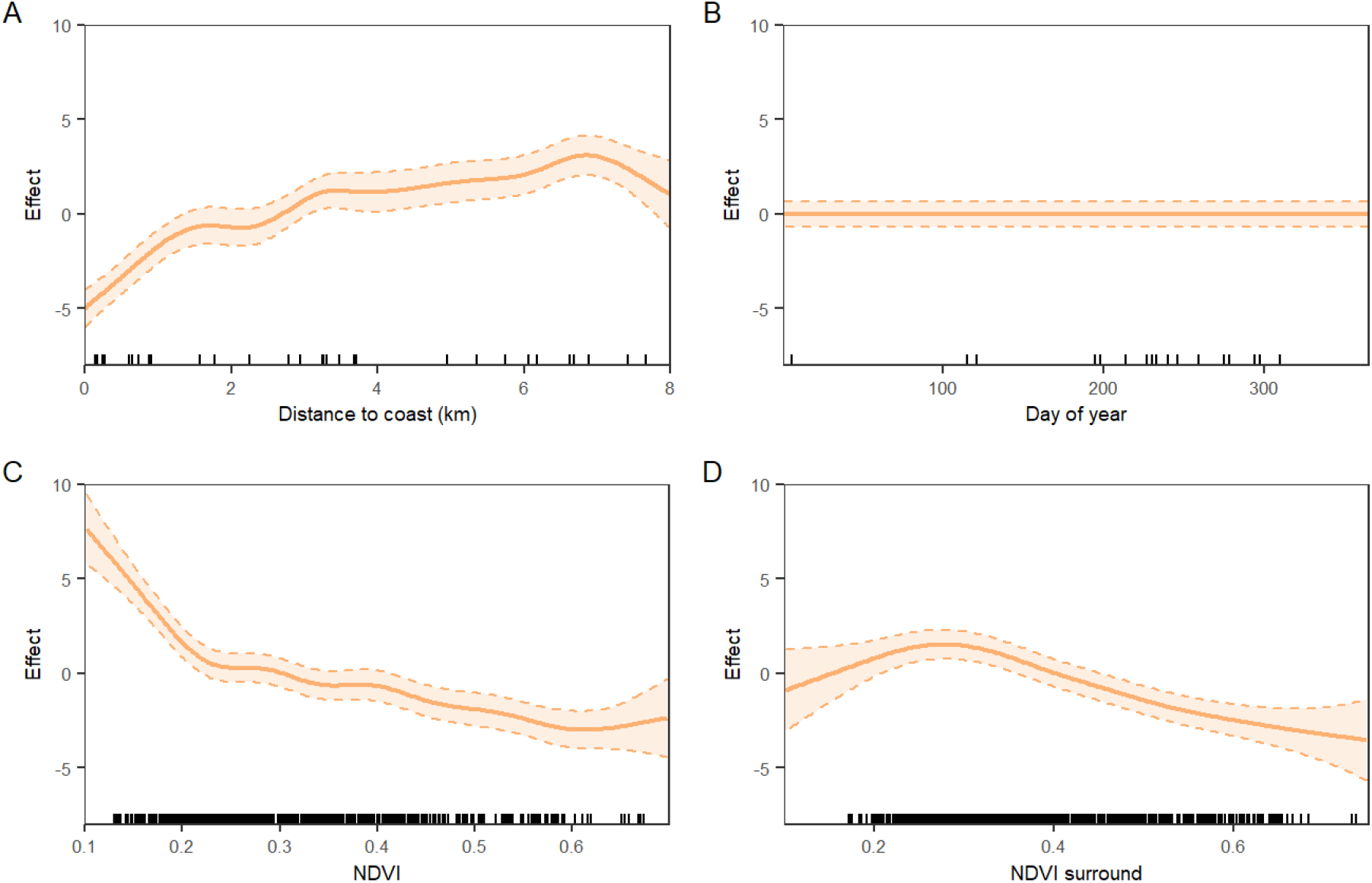
Modelled effects of predictors of surface temperature in informal settlements for A) distance to coast, B) seasonal trend, C) mean NDVI of a settlement and D) mean NDVI within 250 m radius surrounding the settlement. Ticks above the x-axis represent sampling points included in the model.

Satellite-derived surface temperature was most strongly related to mean indoor temperature (R^2^ = 0.413, P = 0.001) in informal settlements, though relationships with indoor minimum, indoor maximum, outdoor maximum and outdoor mean temperature were also positive and significant (Figure 6). In all cases, the slope of the line was greater than one, where small increases in ambient temperature are associated with larger change in surface temperature. Although the relationship is not one to one, this does suggest that satellite-derived UHIs can predict heat stress experienced by residents of informal settlements, under cloud-free conditions, where elevated surface temperature is representative of hot ambient conditions.

**Figure 6.**
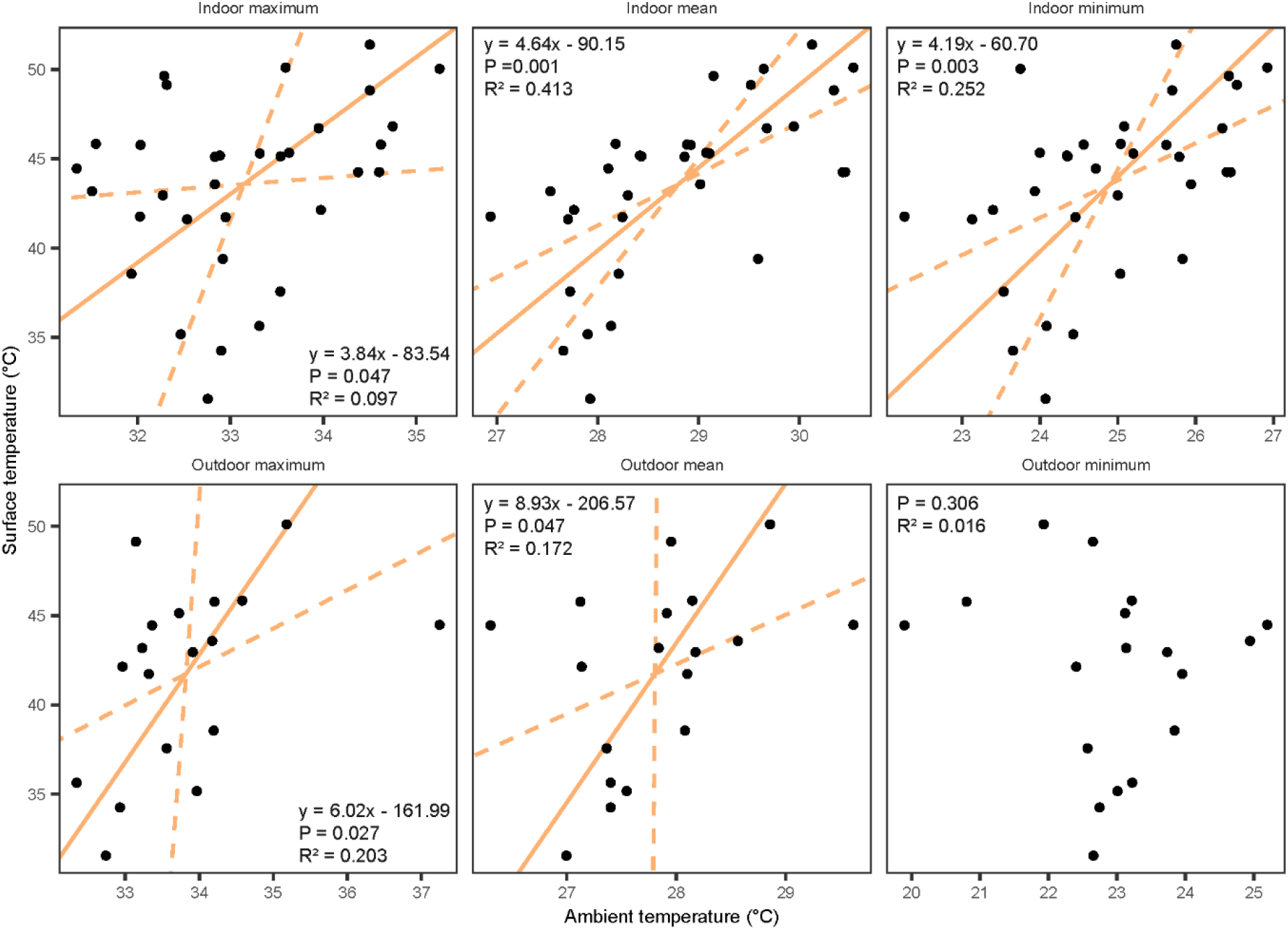
Ranged major axis regression and 95% confidence intervals (dashed) of daily *in situ* ambient temperature variables and satellite-derived surface temperature in a subset of 12 informal settlements with the regression equation, p-values and R^2^. The regression equation is not shown where P > 0.05 and R^2^ close to zero.

## 4. Discussion

Over the past 30 years urban land cover in Makassar has nearly tripled, accompanied by the spatial expansion of UHIs, which affect large parts of the city, including informal settlements. Annual increase in urban land cover of 6.7% exceeded global rates of 3.5% over the same time period, although was less than estimates of 10.8% for Indonesia as a whole, which included larger cities (He *et al*. 2018). Urban heat islands in Makassar averaged 4.5 °C annually in the urban core, and 2.6 °C in informal settlements, although with seasonal fluctuations they reached maxima of 9.2 °C and 6.3 °C above non-urban surface temperature, respectively. These well exceed annual average satellite-derived surface UHIs of 1.3 °C in the tropics (Chakraborty & Lee 2019) or 1.2 °C in Asia (Peng *et al*. 2012), reported by global studies which make simplified comparisons between urban and non-urban pixels within an urban extent. Our results are more in line with surface UHIs between 5 °C and 8 °C detected in South-East Asian megacities, Bangkok, Manila & Ho Chi Minh, during the dry season (Tran *et al*. 2006). Our results highlight that smaller cities, and informal settlements within these, are not exempt from UHIs, and the likely impacts thereof, that have been observed in larger cities. Exposure to elevated temperature has direct effects on health and wellbeing (Mendez-Lazaro *et al*. 2018; Ebi *et al*. 2021b), exacerbates socioeconomic stressors (Tran *et al*. 2013), especially for informal settlement residents, and is expected to continue doing so. The anthropogenic heat burden has already been responsible on average for 37% of warm-season heat-related deaths globally (Vicedo-Cabrera *et al*. 2021).

In contrast to previous studies (Chakraborty *et al*. 2019; Jacobs *et al*. 2019), which identified elevated UHIs in lower income neighbourhoods within cities, we found that UHIs in informal settlements were on average 1.93 °C lower across a year than the urban core (representative of formal settlements), although we could not make direct comparisons to more socioeconomically stable neighbourhoods. Despite the smaller magnitude of UHIs, residents of informal settlements are especially vulnerable to heat due to poor quality housing, a large existing health burden and occupational heat exposure (Tran *et al*. 2013; Ezeh *et al*. 2017). Green space in, and surrounding settlements, along with proximity to the coast appeared to protect informal settlements from the worst UHIs. Green space has a cooling effect both through shading and evapotranspiration (Aflaki *et al*. 2017), although the cooling potential from evapotranspiration is upper bounded by high humidity in the tropics (Yu *et al*. 2018; Manoli *et al*. 2019; Cuthbert *et al*. 2022). Meanwhile, large water bodies, including oceans and lakes, have a cooling effect due to the temperature differential between land and water producing cool onshore breezes during the day (Bonan 2002; Adams & Smith 2014; Cai *et al*. 2018). Settlements with NDVI close to that of the urban core (~0.2) were around 5 °C warmer than settlements with more green space. Settlements within 2 km of the coast also had lower temperatures, with cooling effects of up to 5 °C (Figure 5), although these settlements may be at higher risk of other environmental hazards such as flooding or sea-level rise (Satterthwaite *et al*. 2020). Based on the relationship with mean indoor temperature, surface temperature effect sizes of 5 °C, from proximity to green or blue space, could represent indoor cooling or heating of more than 1 °C.

It is typically thought that UHIs increase in magnitude as a city increases in size and population (Li *et al*. 2017; Zhou *et al*. 2017; Mentaschi *et al*. 2022), though this effect is smaller in tropical cities (Manoli *et al*. 2019). We found that UHIs in long-urbanised areas in Makassar have remained stable since at least the 1990s, in line well-researched cities such as London (Jones & Lister 2009), but have expanded into recently urbanised areas, following patterns observed across Europe (Trusilova *et al*. 2009) and Africa (Li *et al*. 2021). Patterns of UHI development in recently urbanised areas varied with urban land use. Industrial areas unexpectedly declined in surface temperature as patches became urbanised, likely due to the form of industrial buildings which have light coloured, high albedo roofs that reflect heat, and the large size of which can create shade and reduce impervious ground surfaces (Connors *et al*. 2012). By contrast, in other urban patches, which include suburban areas, UHIs develop early on as land becomes urban, with nearly three quarters of surface temperature increase occurring before a patch became 50% urbanised, after which trends stabilised. Given that 70% of new urban land through to 2050 is projected to occur in the tropics (Huang *et al*. 2019), a large portion of which is likely to include informal settlements (Satterthwaite *et al*. 2020), the spatial extent of UHIs is likely to increase substantially. It may, therefore, be preferable to prioritise development, including heat mitigation strategies, of partially urbanised land where UHIs have already developed. Reducing urban sprawl not only limits the spatial expansion of UHIs but also has positive effects for biodiversity by reducing habitat loss (Simkin *et al*. 2022).

Across our results, seasonal patterns of UHIs were driven by increased temperature in September and October in non-urban areas, whilst surface temperature in urban areas is relatively stable year-round (Figure 2; Figure 3). This is in line with previous work suggesting that the magnitude of UHIs is largely controlled by greening in non-urban areas, which increases the temperature differential with urban areas (Yao *et al*. 2019). For example, we observe the largest UHIs in the wet season when non-urban, primarily agricultural, areas are most green (See Figure S2 for monthly plots of non-urban NDVI in Makassar). Towards the end of the dry season, in September and October, however, NDVI declines substantially (Figure S2) as crops are harvested and soil moisture declines (Pandey *et al*. 2014; Kumar *et al*. 2017). Whilst this technically creates an ‘cool island’, relative to the extremely high surface temperatures recorded from predominantly bare earth in the non-urban area (mean 50.7 °C in September and October, from modelled data), surface temperatures in urban areas remain stable and high (mean 46.6 °C in September and October).

Our results highlight the importance of intra-urban analyses to understand spatio-temporal patterns of UHIs and accurately quantify exposure. Mixed long-term trends with urban land use would otherwise be obscured by coarser resolution data or simplified urban – non-urban comparisons. Fine scale data is also essential for understanding UHIs in informal settlements. Even MODIS, one of the most common sources of remote sensed surface temperature data (Kotharkar *et al*. 2018), with a spatial resolution of 1 km^2^ is too coarse to detect patterns across informal settlements which have an average size of 0.016 km^2^ (Friesen *et al*. 2018). Whilst satellite derived surface temperature is not a perfect representation of heat stress experienced by people (Venter *et al*. 2021), we show that it has a reasonable, although not one to one relationship, with *in situ* temperature measured in 12 informal settlements. Impetus is growing for local meteorological monitoring to capture microclimate across complex landscapes, including cities (Zaitchik & Tuholske 2021). Such data can be collected through low-cost sensor networks (as used here) which have been successful in the informal settlement context (Scott *et al*. 2017; Ramsay *et al*. 2021; Van de Walle *et al*. 2022) and can be used to corroborate and supplement remote-sensed data (Venter *et al*. 2020). Indeed, such information is essential to understand the magnitude and mechanisms of UHIs in informal settlement globally, and the interactions they may have with climate warming and changing temperature extremes (Chapman *et al*. 2017).

## 5. Conclusions

If the ambitions of the Sustainable Development Goals and Planetary Health movements are to be met, heat mitigation must be prioritised and targeted where large UHIs intersect with vulnerable communities. Our results demonstrate large UHIs across a socioeconomically diverse tropical city, which emerge rapidly as land is urbanised. Given that UHIs tend to stabilise after land becomes 50% urbanised, the development of partially urbanised areas should be prioritised to spatially constrain UHIs, with consideration of maintaining green space, which mitigates the worst UHIs. Informal settlements, which are often the subject of redevelopment and upgrading programs, represent an opportunity to explicitly consider green space and other heat mitigation strategies such as ventilation and reflective surfaces (Nutkiewicz *et al*. 2018; Kimemia *et al*. 2020). Upgrades, whether nature-based or other, should balance the socioeconomic benefits of built infrastructure, such as roads, with the need to maintain green space. Nature-based solutions offer a compromise where infrastructure upgrades, such as constructed wetlands to improve water and sanitation, can provide co-benefits of heat mitigation by maintaining green and blue space (Wong *et al*. 2020; Leder *et al*. 2021; Hamel & Tan 2022). Such localised interventions will be essential for not only improved local conditions but ongoing climate resilience (Estrada *et al*. 2017), especially given the already high anthropogenic heat mortality burden (Vicedo-Cabrera *et al*. 2021) and forecasts for average global temperature increases of at least 2 °C over the coming decades (Meinshausen *et al*. 2022).

## Acknowledgements

This research was supported by The Wellcome Trust [Our Planet, Our Health Grant 205222/Z/16/Z]. Research was conducted with permission from the Indonesian Ministry of Research and Technology. E.E.R is supported by an Australian Government Research Training Program Scholarship. G.A.D. is the recipient of an Australian Research Council Discovery Early Career Researcher Award (DE190100003) funded by the Australian Government. We thank the RISE Program Consortium (details of the RISE study can be found on the study website: www.rise-program.org) for field data collection assistance and administrative support.

## Ethical statement

Ethics review and approval was provided by participating universities and local IRBs, including Monash University Human Research Ethics Committee (Melbourne, Australia; protocol 9396) and the Ministry of Research, Technology and Higher Education Ethics Committee of Medical Research at the Faculty of Medicine, Universitas Hasanuddin (Makassar, Indonesia; protocol UH18020110). The RISE program is a randomised control trial registered on the Australian New Zealand Clinical Trials Registry (ANZCTR) (Trial ID: ACTRN12618000633280).

## CRediT authorship contribution statement

**Emma E. Ramsay:** conceptualisation; investigation; data curation; methodology; software; writing – original draft. **Grant A. Duffy:** conceptualisation; investigation; data curation; methodology; writing – original draft. **Kerrie Burge:** investigation; writing-review and editing. **Ruzka R. Taruc:** investigation; project administration; writing-review and editing. **Genie M. Fleming:** investigation; data curation; writing-review and editing. **Peter A. Faber:** investigation; writing-review and editing. **Steven L. Chown:** conceptualisation; supervision; methodology; writing – original draft.

## Data availability statement

The data and computer code required to reproduce this work are available at [link to be provided upon publication]. Landsat satellite data are freely available from the United States Geological Survey.

