## Supplemental Information for "Spatio-temporal development of the urban heat island in a socioeconomically diverse tropical city"

### Supplementary information

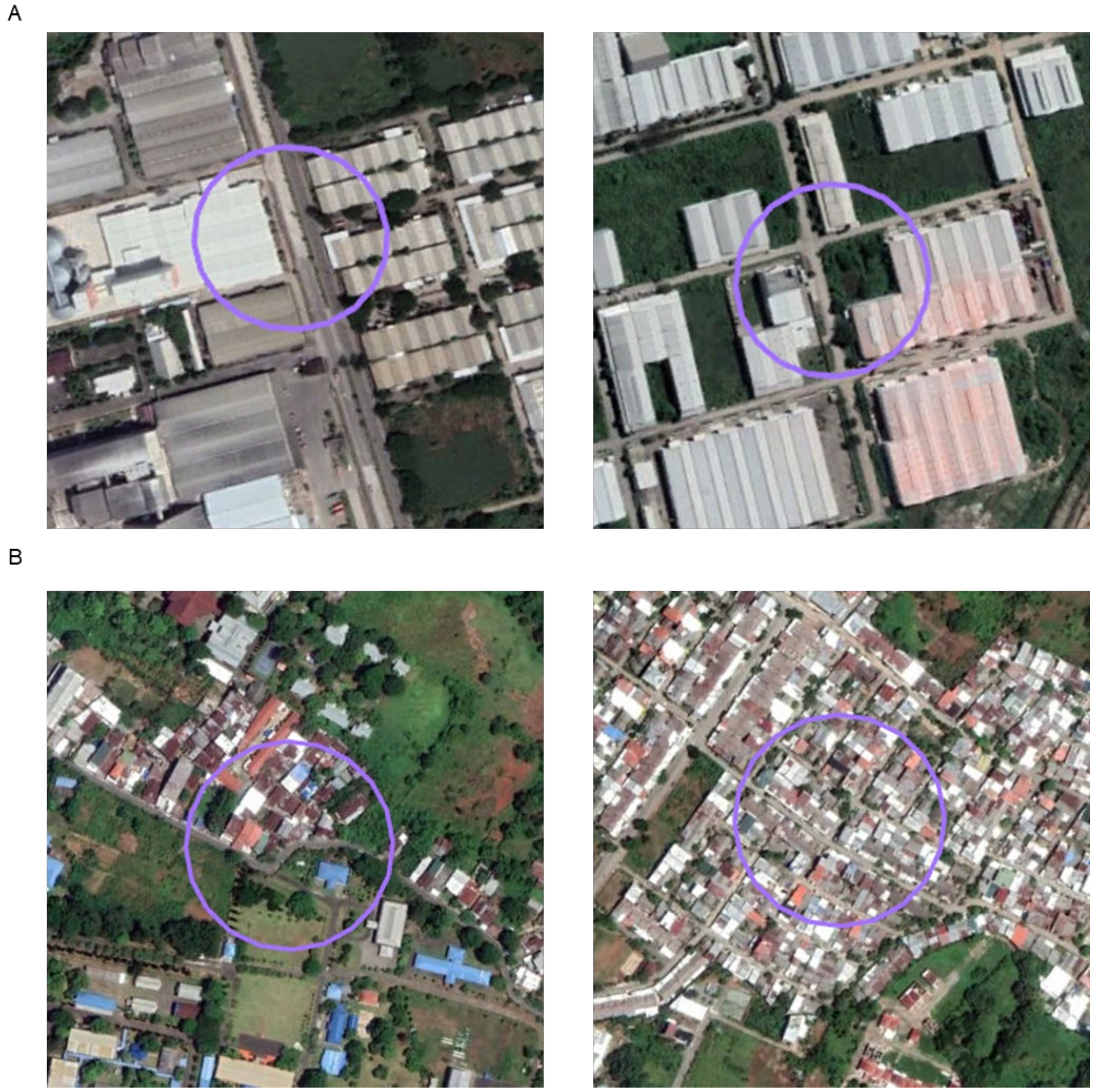

**Figure S1.** Examples of randomly sampled urban change patches classified as A) industrial or B) other urban. Map imagery: Google Earth © 2022 Maxar Technologies.

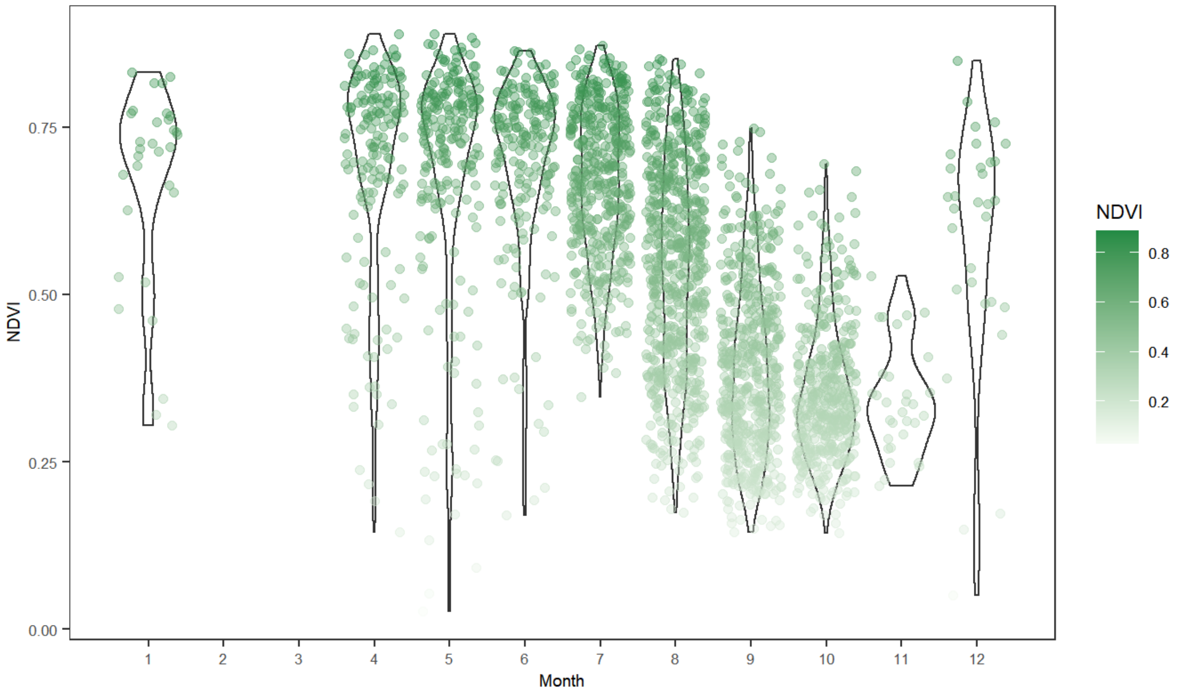


**Figure S2.** Normalised Difference Vegetation Index (NDVI) in randomly sampled non-urban patches by month.

**Table S1.** Summary of *in situ* temperature data collected from loggers in 12 informal settlements on days which overlapped with cloud-free satellite-derived surface temperature. The number of loggers (n loggers) represents the range of loggers in each settlement with data.

| | Outdoor | | Indoor | |
|---|---|---|---|---|
| **Satellite overpass date** | **n settlements with data** | **n loggers** | **n settlements with data** | **n loggers** |
| 6$^{th}$ Jan 2019 | 12 | 3-5 | 12 | 45-60 |
| 17$^{th}$ Jul 2019 | 2 | 3 | 5 | 8-26 |
| 2$^{nd}$ Aug 2019 | 1 | 2 | 2 | 16-23 |
| 18$^{th}$ Aug 2019 | 1 | 2 | 2 | 16-23 |
| 3$^{rd}$ Sep 2019 | 1 | 2 | 2 | 16-23 |
| 5$^{th}$ Oct 2019 | 1 | 2 | 8 | 2-36 |
| 21$^{st}$ Oct 2019 | 0 | 0 | 1 | 5 |
| 6$^{th}$ Nov 2019 | 0 | 0 | 1 | 5 |

**Table S2.** Summary of final General Additive Mixed Models (GAMMs), formulas, time points included in the model (n), adjusted $R^2$ and corresponding figure in the main text. For GAMM formulas $f_n$ are smooth terms, $i$ refers to the temperature observation, $j$ the spatial area (urban core, urban change, informal settlements or non-urban) and $k$ the randomly sampled patch or settlement. All models included residual spatial correlation structures modelled as *corExp* and nested within each time point. Smooths of "days" are long term trends modelled as number of days since first sampling date and "doy" represents the seasonal trend modelled as day of year.

| GAMM | GAMM Formula | n | Adj-R² | Fig |
|---|---|---|---|---|
| Long term trends (1991-2020) in surface temperature in the urban core and non-urban areas | $y_{ij} = \alpha + area_i + f_1(days_i)_j + f_2(doy_i)_j + \varepsilon_i$ | 82 | 0.448 | 3 |
| Long term trends in surface temperature in the urban change and non-urban areas | $y_{ij} = \alpha + area_i + f_1(days_i)_j + f_2(doy_i)_j + f_3(days_i, doy_i)_j + \varepsilon_i$ | 80 | 0.513 | 3 |
| Patch level urban change trends | $y_{ik} = \alpha + f_1(days_i) + f_2(days_i)_k + f_3(doy_i) + \varepsilon_i$ | 82 | 0.382 | 4 |
| Recent (2017-2020) seasonal trends in surface temperature in the urban core, informal settlements and non-urban areas | $y_{ij} = \alpha + area_i + f_1(doy_i)_j + \varepsilon_i$ | 17 | 0.478 | 2 |
| Predictors of surface temperature among informal settlements | $y_i = \alpha + f_1(NDVI_i) + f_2(NDVI\ surround_i) + f_3(Distance\ to\ coast_i) + f_4(doy_i) + \varepsilon_i$ | 17 | 0.385 | 5 |

**Table S3.** Predictors of surface temperature among informal settlements modelled as linear parametric terms in a GAMM with a seasonal smooth. Adjusted- $R^2 = 0.35$. edf = effective degrees of freedom.

| **Predictor** | **Estimate** | **SE** | **t** | **P** |
|---|---|---|---|---|
| NDVI | -10.58 | 1.05 | -10.00 | < 0.001 |
| NDVI surround | -11.02 | 1.11 | -9.92 | < 0.001 |
| Distance to coast (km) | 0.85 | 0.09 | 9.23 | < 0.001 |
| **Smooth terms** | **edf** | | **F** | **P** |
| Day of year | 4.623 | | 2.026 | 0.003 |
